# 3.7 Å cryo-EM structure of the core Centromere Binding Factor 3 complex

**DOI:** 10.1101/241067

**Authors:** Wenjuan Zhang, Natalya Lukoynova, Shomon Miah, Cara K Vaughan

## Abstract

The Centromere Binding Factor 3 (CBF3) complex binds the third Centromere DNA Element in organisms with point centromeres, such as *S. cerevisiae*. It is the only essential centromere binding complex as it facilitates genetic specification of point centromeres. It is therefore the most fundamental complex of the kinetochore in these organisms and its association with centromere DNA allows association of all other kinetochore components. We have determined the atomic structure of the core complex of CBF3, comprising 3 of its 4 components, using cryo-EM. The architecture of the complex is 'U'-shaped, with a deep, strongly basic channel that is narrow at one end and wide at the other. Combining our structure and in vitro assays, we present a model for its association with centromere DNA.

## Introduction

The integrity of genetic information passed through generations relies on faithful segregation of chromosomes during mitosis. The kinetochore, a mega-Dalton protein assembly, enables this segregation by specifically associating with both the centromere (CEN) of sister chromatids and the microtubules of the mitotic spindle. Most eukaryotes have regional centromeres with unique satellite repeat structures that vary in length but whose arrangement is largely conserved between chromosomes despite being unconserved in sequence. Regional centromeres are specified epigenetically, by the presence of an essential centromeric histone H3 variant, CENP-A, which is found at CEN DNA, where it is interspersed with canonical nucleosomes (Verdaasdonk and Bloom, 2011).

By contrast budding yeasts, including *S. cerevisiae*, have evolved point centromeres, comprising conserved, short and essential CEN DNA, comprising three Centromere DNA Elements (CDEs), typically of ~125 bp. Point centromeres evolved from an ancestor with an epigenetically-specified centromere (Malik and Henikoff, 2009) and this evolutionary transition introduced genetic specification of the centromere while retaining aspects of epigenetic specification, in particular the essential presence of the Cenp-A homologue, Cse4.

Unique to these organisms, the Centromere Binding Factor 3 (CBF3) complex provides a physical link between the genetic and epigenetic mechanisms of centromere specification. It associates specifically to the highly conserved CDEIII (Lechner and Carbon, 1991) and is responsible for deposition of Cse4, through a direct interaction with the Cse4 chaperone, Scm3/HJURP (Camahort et al., 2007). Its epigenetic role is emphasized by the observation that a synthetic kinetochore can be assembled in the absence of centromere DNA elements provided functional CBF3 is available to stabilise Cse4 incorporation at the centromere (Ho et al., 2014).

CBF3 comprises two homodimers, of Cep3 and Ndc10, and a Ctf13-Skp1 heterodimer. Cep3 provides sequence specificity through binuclear zinc-cluster domains homologous to those found in GAL4-like fungal transcription factors (Lechner, 1994; MacPherson et al., 2006). These domains bind a pseudo TGT/CCG palindrome in CDEIII (Espelin et al., 1997). Ndc10 contributes both non-specific DNA-binding (Cho and Harrison, 2011; Espelin et al., 1997) and association with the centromeric histone chaperone, Scm3/HJURP (Camahort et al., 2007). The Skp1-Ctf13 heterodimer interacts with both Cep3 and Ndc10 and interacts with CDEIII at a completely conserved G centrally positioned between the binding sites for the binuclear zinc-cluster domains (Espelin et al., 1997).

The molecular mechanism of CBF3 association with CDEIII is unknown, despite a wealth of genetic and biochemical data and crystal structures of domains from individual components (Bellizzi et al., 2007; Cho and Harrison, 2011; Perriches and Singleton, 2012; Purvis and Singleton, 2007), due to an absence of atomic-resolution structural information for the CBF3 complex as a whole.

Herein we present the cryo-EM structure of the CBF3 core complex (CBF3CCΔN) at atomic resolution. The complex forms a deep channel that is both highly charged and strongly conserved, and is perfectly sized to accommodate DNA. The structure identifies structural elements from Ctf13 that contribute to DNA association and our in vitro experiments confirm this role and show that this association is sequence independent. The data allow us to present a model, which accounts for previously published cross-linking experiments and provides a high resolution view of the CBF3CCΔN-CDEIII structure.

## Results

### Cryo-EM Studies of CBF3

Dissection of the assembly and turnover of CBF3 *in vivo* indicated that Cep3 and Ctf13 associate early in the CBF3 assembly pathway while association with Ndc10 occurred at much later time points (Rodrigo-Brenni et al., 2004). These results were in agreement with earlier *in vitro* reconstitution experiments (Russell et al., 1999) therefore we expressed and purified a CBF3 core complex (CBF3CC) comprising the Ctf13-Skp1 heterodimer and the Cep3 homodimer (Figure 1A). Full length Cep3 rendered CBF3CC unstable, but the complex with an N-terminally truncated Cep3, in which the binuclear zinc cluster domains were missing, yielded a 220 kDa complex (CBF3CCΔN) that was purified to homogeneity (Figure S1A) and was suitable for cryo-EM studies (Figure S1B). 2D-classification of motion-corrected cryo-EM images generated classes with clear secondary structural details (Figure S1C) that enabled *de novo* reconstruction of a 3D map. After refinement, the final 3.7 Å map comprised 187606 particles with good angular distribution (Figure S1D), judged using the FSC gold standard method (Figure S1E). The final map has a horseshoe shape with a deep central channel (Figure 1B).

**Figure 1:**
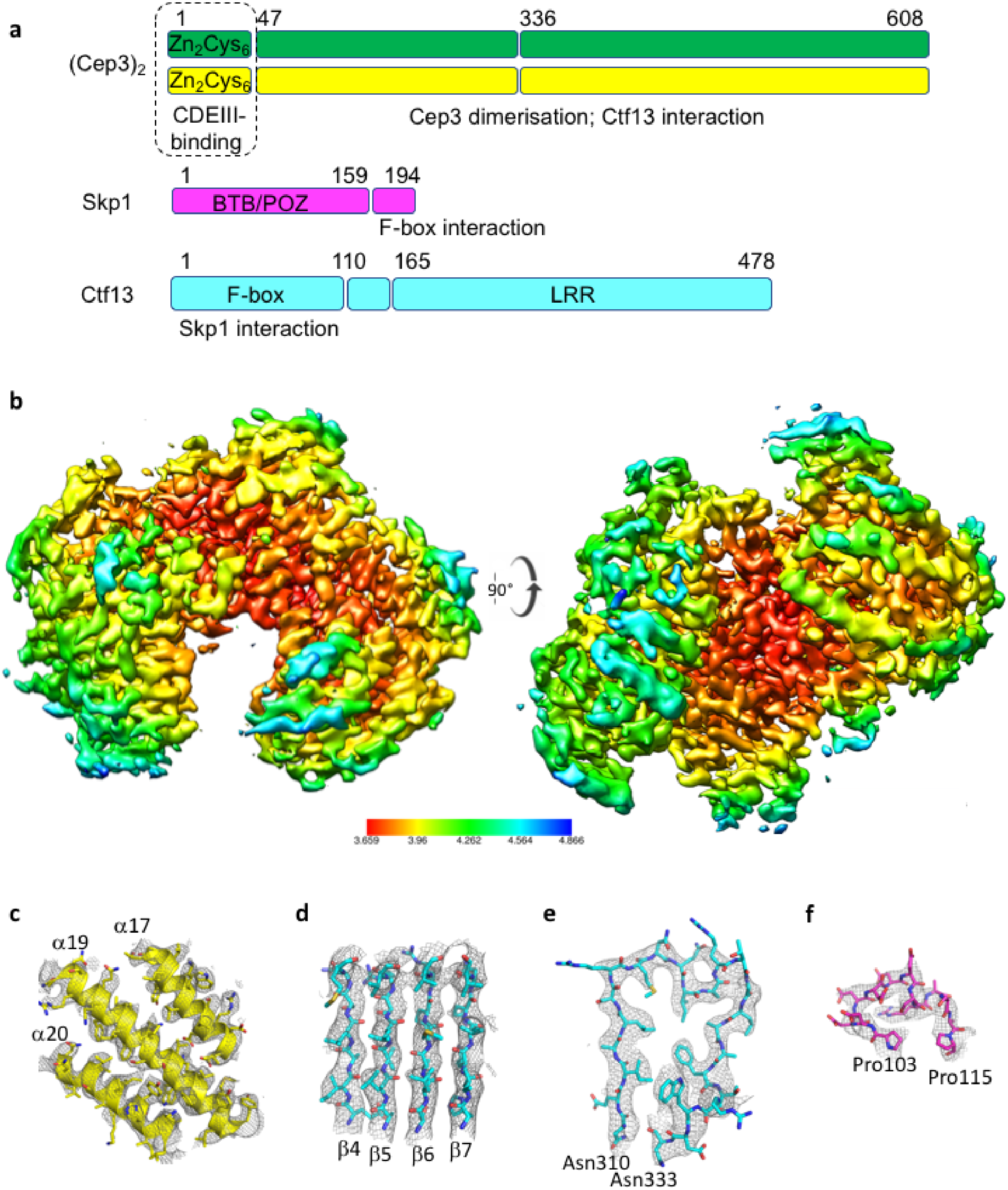
Cryo-EM reconstruction of the CBF3CCΔN. (A) Components of CBF3CCΔN, annotated with domain boundaries and architecture. Known binding-partners are indicated below the relevant domains. Domains not included in the construct used for structure determination are boxed with a dotted line. (B) Structure of CBF3CCΔN coloured by local resolution. The overall resolution estimate is 3.7 Å. Representative electron density for (C) helices from Cep3ΔN, (D) 4 beta-strands of the LRR beta-sheet from Ctf13, (E) LRR4 from Ctf13, (F) an acidic loop in Skp1.

### Atomic Model of CBF3CCΔN

Secondary structure details and side chains were clearly visible throughout the map (Figures 1C-F) enabling an atomic model of the structure to be built and refined (Figures 2A-D, S1F). The Cep3 homodimer was readily recognised, facilitated by its largely helical structure and the identification of an approximately two-fold axis, that could also be seen in the 2D classes (Figure S1C). The 2.5 Å crystal structure of the Cep3 homodimer (Purvis and Singleton, 2007) was fit in the map as a rigid body and this structure is essentially unchanged after refinement with a final Ca RSMD of 1.1 Å. Of the remaining density, the most striking secondary structural feature is an 8-stranded parallel beta sheet. This is part of a larger solenoid structure, corresponding to the predicted LRR fold of Ctf13, and comprises 8 LRR motifs (Figure S2A). The LRR domain is decorated at each end by additional domains (Figures 2B, S2A).

**Figure 2:**
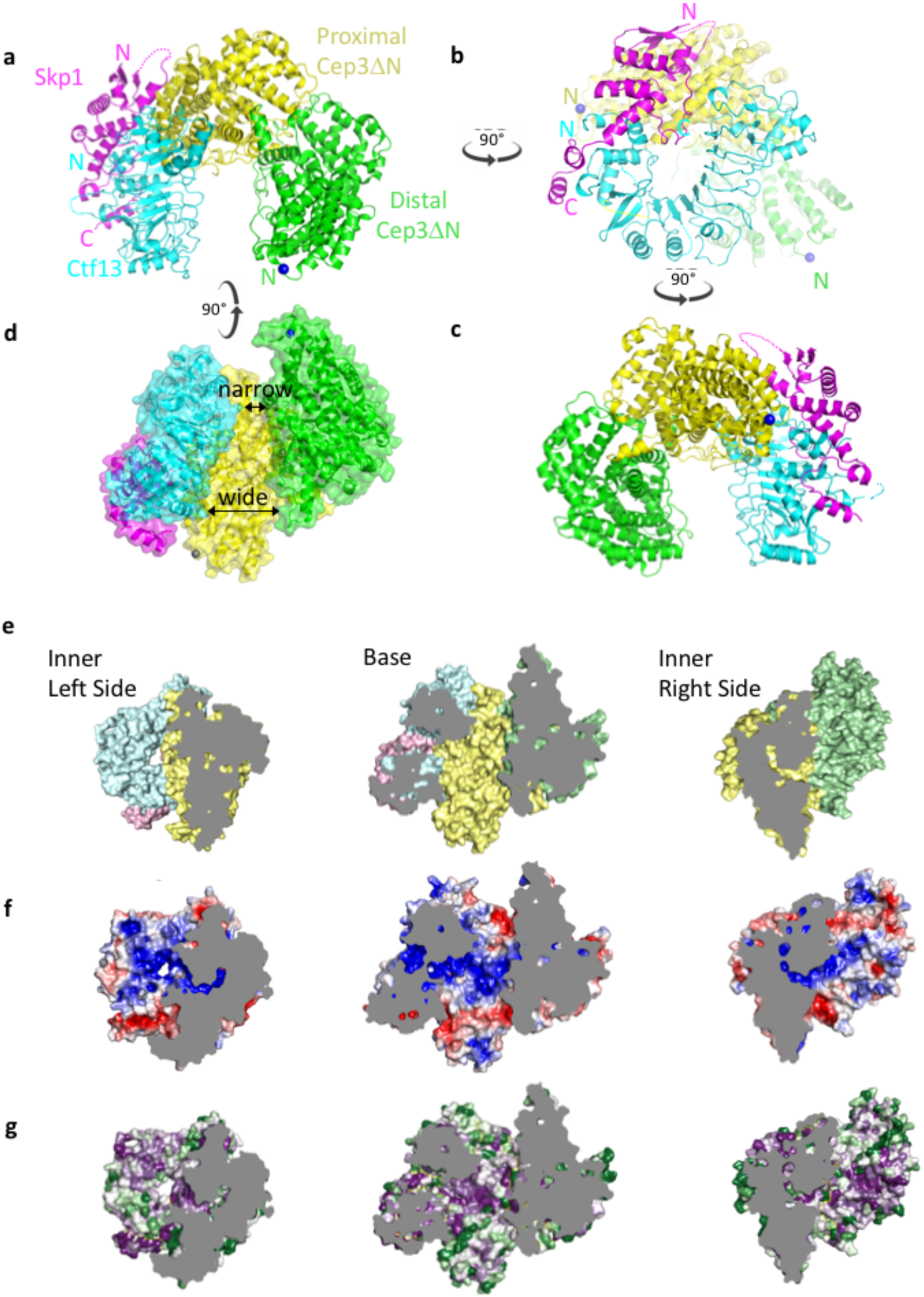
Atomic model of CBF3CCΔN. (A-D) The structure of CBF3CCΔN with Ctf13 (cyan), Skpl(pink) and the Cep3ΔN homodimer (yellow and green) showing 4 views each related by 90°. One view (D) highlights variation in the width of the channel. The N-termini of the Cep3ΔN homodimer are shown as blue balls. Views of each side of the inner surface of the channel coloured by (E) protein (colour as A-D above), (F) electrostatic potential (from −5 (red) to +5 (blue) TeV) and (G) by conservation (purple-white-green = high-medium-low conservation).

Into one of these, the N-terminal BTB/POZ domain of yeast Skp1 (Orlicky et al., 2003) could be fit as a rigid body. The map included density for an acidic loop that is rarely visible in Skp1 crystal structures (Figure 1F). In typical Skp1-Fbox interactions, the ~30 residues after the BTB/POZ domain form 2 C-terminal helices (a7 & a8) that wrap closely around the F-box of the partner protein, forming a compact structural domain that is conserved in all published Skp1-Fbox structures to date. However, this canonical Skp1-Fbox structure does not fit in the CBF3CCΔN map. The position of the first helix of the F-box of Ctf13 is not compatible with the common orientation of Skp1-α7 and consequently both α7 & α8 of Skp1 are reoriented by 86° and 60° respectively (Figures S2B, S2C, S3A, S3B). The F-box is connected to the LRR domain by a linker subdomain comprising a 3-stranded anti-parallel β-sheet and a long α-helix (Figure S2A). The majority of the F-box of Ctf13 is well conserved between Saccharomycetaceae, therefore this atypical structure is likely to be found in all Skp1-Ctf13 homologues. In addition, there is an ~50 amino acid insertion between α2 and α3 of the F-box for which there is no electron density. Since some of this has weak conservation, it may be structured in the presence of other binding partners.

The remaining density is an α-β subdomain that decorates the C-terminal end of the Ctf13 LRR, formed by insertions within the last 3 LRRs of Ctf13. This subdomain contacts the previously mentioned acidic loop of Skp1 such that, overall, the Skp1-Ctf13 heterodimer forms a toroidal structure in which both the N-terminal F-box and the C-terminal α-β LRR-insertion domain of Ctf13 associate with either end of Skp1 (Figures 2B, S3C). Other LRR-containing F-box proteins form toroids, notably both the TIR1 and COI1 plant hormone receptors (Sheard et al., 2010; Tan et al., 2007), but in these cases the toroid is formed by the LRR domain alone, with Skp1 and the LRR domains almost perpendicular to each other. By contrast for Skp1-Ctf13, the unusual Skp1 – F-box interaction forces both Skp1 and Ctf13 into the same plane (Figures 2B, S2D).

The Ctf13-Skp1 heterodimer forms the left side of the horseshoe, and makes extensive contacts with the base, formed by the 'proximal' monomer of the Cep3 homodimer (Figures 2A-D, S3D-F). This interface includes density for part of a loop in Cep3, from residues 330 to 339 not previously visible in the crystal structures. The 'distal' monomer of the Cep3 dimer forms the remaining side of the horseshoe (Figures 2A-D). This arrangement positions the (Cep3ΔN)_2_ N-termini, and consequently the truncated binuclear zinc cluster domains, at opposite ends of the channel created by the horseshoe. The position of Ctf13 relative to (Cep3ΔN)_2_ results in the channel being considerably narrower next to the N-terminus of the distal Cep3ΔN monomer (Figure 2D).

### CBF3CCΔN channel is the putative binding site for CDEIII

Ctf13 and Cep3 line the channel with basic residues that are strongly conserved between Saccharomycetaceae (Figures 2E-G). In Ctf13 a series of arginine and lysine residues extend like fingers from the inter LRR turns of LRRs 1–6 into this groove (Figure S2E). LRR3 projects two neighbouring arginines, Arg307 and Arg308, and the latter is positioned directly along the twofold axis of (Cep3ΔN)_2_, at the midpoint between the truncated (Cep3ΔN)_2_ N-termini (Figure 3A). Additional conserved basic residues from Cep3ΔN extend towards the channel from each Cep3 protomer, including Lys265, Arg273 and Lys364.

**Figure 3:**
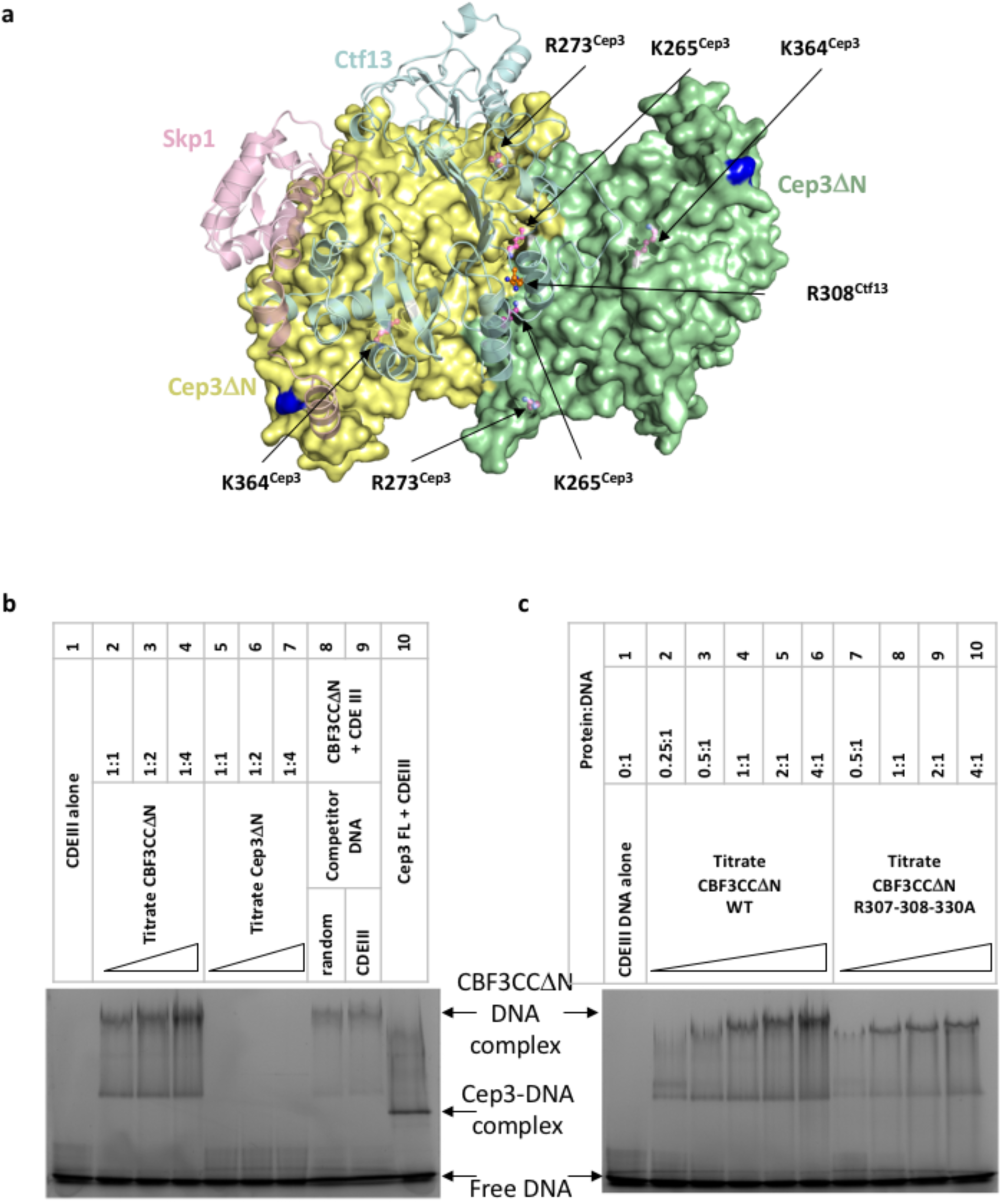
CBF3CCΔN channel is the putative binding site for CDEIII. (A) View down the two-fold axis of the Cep3ΔN homodimer. Cep3ΔN dimer surface is coloured yellow & green; Skp1 (pink) and Ctf13 (cyan) are semi-transparent and shown as cartoon. R308^Ctf13^ lies directly along the two-fold axis (orange ball and stick). Conserved basic residues in Cep3 are highlighted in magenta ball and stick. (B) Electrophoretic mobility shift assays performed with (A) 1.6 μM fluorescently-labelled CDEIII DNA and a titration of CBF3CCΔN (lanes 2–4) or Cep3ΔN (lanes 5–7) or 6.4 μM of full length Cep3 (lane 10). Lanes 8 and 9 show competition of CBF3CCΔN binding with 80 μM unlabelled probe. (C) 1.6 μM fluorescently-labelled CDEIII DNA with a titration of CBF3CCΔN (lanes 2–6) and CBF3CCΔN with Ctf13 mutations R307A-R308A-R330A.

The charge and conservation within the channel as a whole, and the striking relative orientation of the arginine residues from neighbouring LRRs in Ctf13, suggested that the channel may provide the binding site for CEN DNA, with these residues potentially contributing direct interactions with CDEIII. In order to test this model, we carried out electrophoretic mobility shift assays. Full length Cep3 binds CDEIII tightly whereas a construct of Cep3 in which the N-terminal binuclear zinc cluster domains are truncated, Cep3ΔN, does not bind CDEIII DNA, consistent with previous observations (Figure 3B; compare lanes 5–7 and 10) (Purvis and Singleton, 2007). By contrast CBF3CCΔN shows a DNA-gel shift (Figure 3B; compare lanes 2–4 with lanes 5–7) indicating that association with the Skp1-Ctf13 heterodimer significantly enhances the affinity of Cep3ΔN for CDEIII DNA. Since CBF3CCΔN has the binuclear zinc cluster domains of Cep3 truncated, we tested whether the association was sequence-specific. Labelled CDEIII DNA could be competed with either unlabelled CDEIII DNA or a random DNA duplex of equal length (Figure 3B; lanes 8&9). Our in vitro data therefore provide conclusive evidence that the binuclear zinc cluster domains are not required for the association of the CBF3CC complex with DNA. However the binuclear zinc cluster domains from Cep3 are the sole determinants of sequence specificity, as the remaining component of the full CBF3 complex, Ndc10, has also been shown to contribute affinity not but specificity to the CDEIII association (Cho and Harrison, 2011).

We atested the contribution of the highly conserved Arg307, Arg308 and Arg330 from Ctf13 to this interaction. The triple mutation to alanine reduces association of CBF3CCΔN with DNA, consistent with a model in which these residues contribute affinity to DNA-binding (Figure 3C).

## Discussion

### Model for CBF3CC association with CDEIII

Our results support a model in which CDEIII DNA binds in the deep channel of the CBF3 core complex. Consistent with this the diameter of the channel accommodates a DNA duplex (Figure 4A).

**Figure 4:**
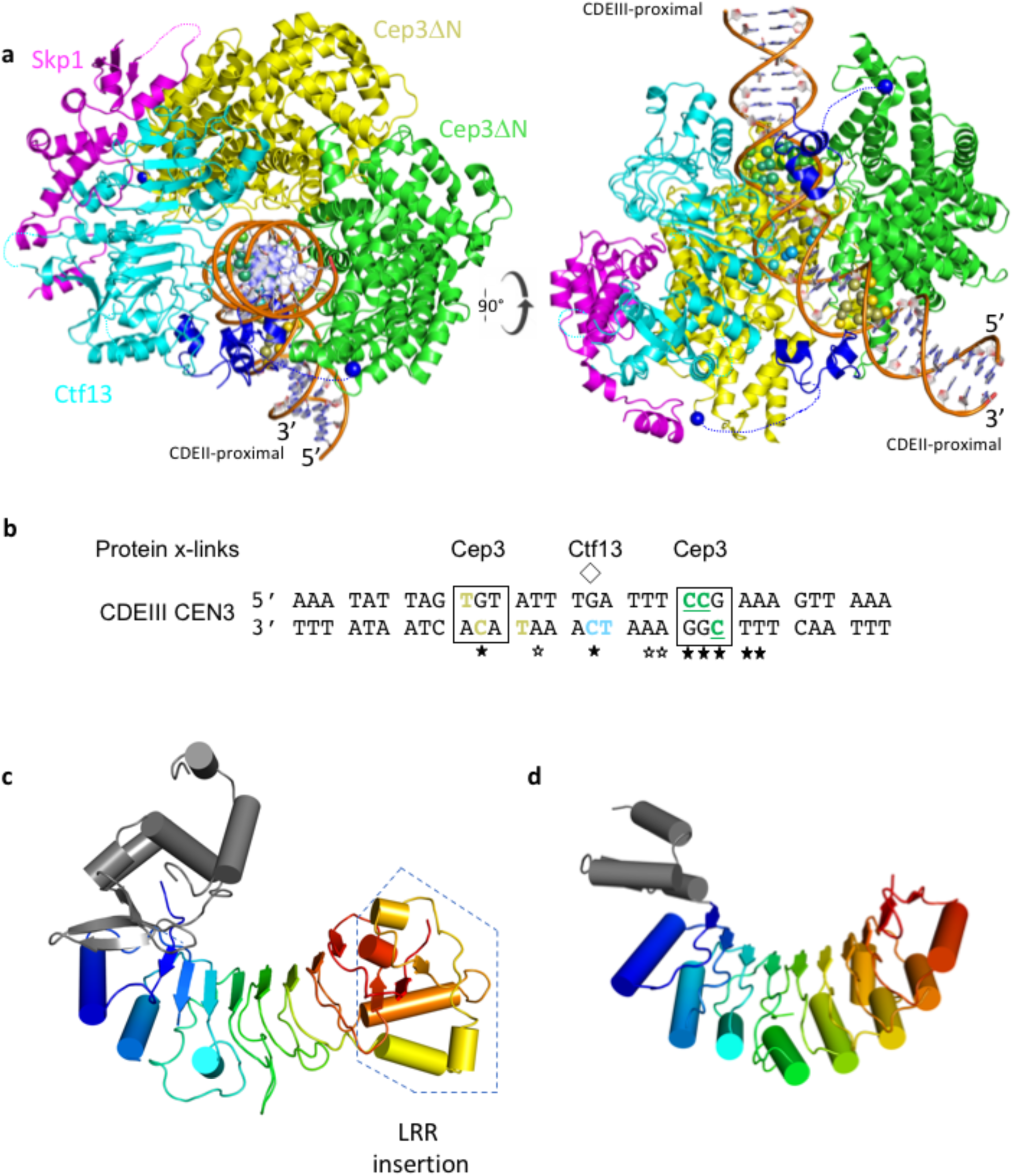
CBF3CCΔN model bound to CDEIII and structural homology of Ctfl3. (A) Model of CBF3CCΔN with CEN 3-CDEI11 DNA bound in the channel. DNA was modelled with a 55° bendbetween the half-sites using 3D-DART (van Dijk and Bonvin, 2009). The CCG half-site is coloured green, the central conserved G is coloured cyan, and the TGT half-site is coloured yellow and their locations emphasized using ball representation for the ribose and base. The end closest to CDEII is labelled. The Cep3 binuclear zinc cluster domains (dark blue) are modelled by superposition of the Gal4-CCG structure (Marmorstein et al., 1992) (PDB 1d66) on each halfsite. (B) The sequence of CEN3 CDEIII. Half-sites of the pseudo-palindrome are boxed. The pseudo-dyad axis is marked with a diamond. Completely conserved and strongly conserved bases (15 of 16 chromosomes) are indicated with filled or empty stars respectively. Bases that interact with Cep3 or Ctf13 are highlighted in colour; bases whose labelling interfered with CBF3 binding are underlined. Cartoon representations of (C) Ctf13 and its closest structural F-box containing homologue (D) human KDM2B (PDB 5jh5). The LRR domains are coloured rainbow and the F-box is coloured grey.

Previous crosslinking data identified crosslinks between Ctf13 and the completely conserved cytosine at the pseudo-dyad axis, and its neighbouring 3' thymine, on the bottom strand of CEN3 CDEIII (Espelin et al., 1997) (Figure 4B). Alignment of the pseudo-dyad axis of modelled CEN3 CDEIII DNA with the twofold axis of Cep3 places the central cytosine in line with the conserved Arg308 of Ctf13. If the DNA is then oriented such that the most conserved surface of Ctf13 aligns with the conserved 3' end of CDEIII, the CCG half-site is placed at the narrow point of the channel (Figure S4A). There is strong sequence conservation between Cep3 and the binuclear zinc cluster domains of fungal transcription factors with known structure, including between residues that contribute to half-site recognition (Figure S4D). If the binuclear zinc cluster of Cep3 is modelled using the prototypical GAL4 crystal structure (Marmorstein et al., 1992), the narrow gap perfectly accommodates the binuclear zinc cluster domain and orients its C-terminal end towards the N-terminal end of Cep3ΔN (Figure S4B). This superposition packs the binuclear zinc cluster domain against a strongly conserved loop between the 3^rd^ and 4^th^ LRR motifs. The equivalent superposition at the other half-site is sterically incompatible (Figure S4C). However, modelling a bend in CDEIII of 55°, as observed in AFM images of the CBF3 complex bound to CEN DNA, allows the second binuclear zinc cluster domain to be readily accommodated (Figure 4A). This model places the conserved CCG and TGT half-sites in very different environments: the former is buried by Ctf13 and Cep3 at the narrow end of the channel, while the latter is solvent accessible at the open end of the channel. This wide channel entrance at the TGT half-site suggests that the exact orientation of the bent DNA within the channel is uncertain, however the presence of other factors, such as Ndc10, is likely to sterically lock a single conformation.

In a recently published similar structure of the CBF3 core complex (Leber et al., 2017), in which the fold of the Ctf13 component could be partially assigned, but not its sequence, this second binuclear domain packs in a groove between Ctf13 and Cep3 in a manner that would not allow binding to the TGT half-site. Since this is not compatible with the observation of crosslinks between the TGT half-site and Cep3 (Espelin et al., 1997), we suggest that this could be an inactive conformation for the binuclear zinc cluster domain, and that the presence of DNA may unlock it from the docked position.

Our structure and model explain a wealth of data published over several decades: The structural asymmetry accounts for observed asymmetric chromosome nondisjunction rates seen when CDEIII is altered: mutations centred around the CCG half-site cause rates of chromosome loss up to 2 orders of magnitude greater than mutations of the TGT (Hegemann et al., 1988). Similarly, labelling of the CCG bases is observed to cause significant loss of association with CBF3 compared with the TGT half-site (Espelin et al., 1997), and genetic results identify the CCG triplet as the only bases within the CEN whose substitution cannot be supported in *S. cerevisiae* (Cumberledge and Carbon, 1987; Gaudet and Fitzgerald-Hayes, 1987; Jehn et al., 1991; McGrew et al., 1986; Ng and Carbon, 1987; Niedenthal et al., 1991).

Previous data suggest that the remaining subunit of the CBF3 complex, Ndc10, binds Ctf13 through its N-terminal domain (Cho and Harrison, 2011; Russell et al., 1999). In addition to highlighting the putative DNA-binding surface within the channel, mapping of conservation onto the surface of CBF3CCΔN indicates a strongly conserved contiguous surface on the outside of the horseshoe, comprising Skp1 and the structural elements of the expanded Ctf13 F-box, in particular the 3 stranded antiparallel β-sheet and the fifth α-helix (Figure S4E). If Ndc10 were to bind at this surface it would be close to the CDEII proximal end of CDEIII, potentially placing it in a suitable location for association with Cbf1, a non-essential CDEI-binding protein which has been shown to associate with the N-terminal domains of Ndc10 (Cho and Harrison, 2011). Although crosslinking data suggest that Ndc10 binds to the CDEII distal end of CDEIII (Espelin et al., 1997)1, Ndc10 dimerisation could account for the 'spreading' of Ndc10-DNA contacts to sites downstream of the CCG half-site.

### Structural homology of Ctf13 suggests an evolutionary link to epigenetic modifications of the point centromere

A search of PDBeFOLD and DALI identifies the histone demethylase KDM2B as the closest known F-box containing structural homologue of Ctf13 (Figures 4C-D).

KDM2B is a human lysine demethylase from the JHDM1 family containing the Jumonji (JmjC) domain. In general, this family are responsible for inducing transcriptional silencing through demethylation of H3K36. In lower eukaryotes family members, such as Jhd1 in *S. cerevisiae*, comprise only the histone lysine demethylation domain while in higher eukaryotes they have several additional domains that detect or alter epigenetic states (Klose et al., 2006). In humans these include a zinc finger domain that recognises methylated DNA, and an F-box and LRR domain that can recruit SCF E3 ubiquitin ligase activity (Han et al., 2016; Wong et al., 2016).

Centromeres have been subject to rapid recent evolutionary change, accounting for the wide diversity in centromere sequences between species. Evidence suggests that the point centromeres of Saccharomycetaceae evolved from an ancestor with a regional centromere and the CBF3 complex co-evolved to meet the requirements of genetic specification (Malik and Henikoff, 2009). The structural homology between Ctf13 and KDM2B may indicate an evolutionary path for the Skp1-Ctf13 component of CBF3 in budding yeasts, involving the partition of a KDM2B-like chromatin-associated enzyme in a common ancestor into two independent genes - the JHMD1 family member, Jhd1, and point centromere-associated Ctf13. Whether there is a genetic or physical link between the two resultant genes in extant budding yeasts remains to be determined. Jhd1 has been shown to counter Set2 methylation (Fang et al., 2007), which is associated with RNA pol II transcription (Kim and Buratowski, 2007; Kwon and Ahn, 2011; Sein et al., 2015) and suppression of histone exchange (Venkatesh et al., 2012), but to date, no centromeric function has been attributed. By contrast in S. pombe, the KDM2B-homologue Epe1 contributes to centromere function through regulating heterochromatin boundaries (Trewick et al., 2007). Epe1 may therefore represent an evolutionary intermediate since it functions to define a regional centromere but, like Jhd1, it is a 'minimal' JHDM1 family member, comprising only the JmjC domain without additional targeting domains.

While structural homology alone is not sufficient to indicate functional analogy, it is tempting to speculate that the structural homology of Ctf13 with KDM2B may indicate a molecular link between Ctf13 and epigenetic specification of point centromeres. Such a link has already been shown to exist for the Ndc10 component of CBF3: Ndc10, the remaining component of CBF3 that is not present in our current structure, associates with the histone chaperone Scm3/HJURP, thereby recruiting the centromeric histone Cse4 to the point centromere. Ctf13 may therefore also contribute a currently uncharacterised function in epigenetic specification of the budding yeast point centromere.

## Experimental Procedures

### Protein expression and purification

The yeast expression system was a gift from Dr. Kiyoshi Nagai. His-tagged Cep3 and untagged Ctf13 genes were a gift from Martin Singleton. Our experiments showed that the co-expression of Sgt1 helps the formation and stability of CBF3CCΔN complex therefore we cloned and co-expressed Sgt1 with CBF3CCΔN components. The Sgt1 gene with a C-terminal Strep tag was subcloned into the modified pRS424 vector containing the Ctf13 gene with a C terminal CBP tag. The Skp1 gene was subcloned into the modified pRS426 vector containing Cep3ΔN (47–608) with a N-terminal His tag. Mutants were generated by site-directed mutagenesis PCR. Both plasmids were co-transformed into *S. cerevisiae* yeast strain BCY123 (MATα pep4::HIS3 prb1::LEU2 bar1::HIS6 lys2::GAL1/10GAL4 can1 ade2 trp1 ura3 his3 leu23,112) by using -Trp, - Ura selection plates (Yeast Nitrogen Base, Trp and Ura dropout mix (Formedium Ltd., UK), 55 mg/ml adenine, 55 mg/ml L-tyrosine, 2% glucose). Expression of the complex and mutants were performed in BCY123, as reported. The cells were pre-cultured for 24 hours in selective media and then inoculated into 24 liter non-selective medium with 2% glucose replaced by 2% raffinose to a starting OD of 0.25. The expression of complex was induced with 2% galactose for 16 hours at an OD of 0.9–1.0. Pelleted cells were resuspended in 0.3 times the cell volume in lysis buffer A (50 mM Tris, 500 mM NaCl, 2 mM MgAcetate, 2 mM Imidazole, 4 mM CaCl_2_, 0.2% Igepal CA-630) supplemented with cOmplete™ EDTA-free Protease Inhibitor Cocktail (Roche) and frozen as pellets in liquid N_2_. The pellets were lysed using a freezer miller (SPEX Sample Prep). The complex was purified by Calmodulin resin and eluted by buffer B (10 mM Tris, pH8.0, 500 mM NaCl, 1 mM MgAcetate, 1 mM Imidazole, 4 mM EGTA, 2 mM DTT). Purified fractions were pooled together and loaded onto 1 ml His-Trap column (GE Heathcare), which was pre-equilibrated with buffer C (20 mM Tris, 500 mM NaCl, 20 mM Imidazole, pH8.0, 10 mM β-mercaptoethanol). The eluted protein was concentrated by centrifugal ultrafiltration (Amicon Ultra-15, 10 kDa MWCO, Millipore) and loaded onto Mono Q column (GE Heathcare) after dilution with low salt buffer D (20 mM Tris, pH8.0, 100 mM NaCl, 2 mM DTT, 5 mM EDTA, 10% glycerol). The complex fractions were pooled and concentrated before loading onto a Superdex 200 5/150 GL (GE Healthcare) pre-equilibrated with S200 buffer (15 mM Tris, 200 mM NaCl, 2 mM DTT). The peak fraction was used to make EM grids.

### Cryo-Electron microscopy

The sample (0.12 mg/ml for first dataset of CBF3CCΔN complex, 0.24 mg/ml for second dataset of CBF3CCΔN complex) was applied to glow-discharged Quantifoil 1.2/1.3 300 mesh grids (Agar Scientific). Cryo-EM data was acquired on a FEI Titan Krios at 300 keV, equipped with a K2 Summit direct detector and a GIF Quantum energy filter (Gatan). Data collection was automatically carried out using EPU software (FEI) to record 1236 movies with a defocus range of −1.6 μm to −3.6 μm for the first CBF3CCΔN dataset and 1101 movies with a defocus range of −1.0 μm to −4.0 μm at a magnification of 47170 (1.06 Å pixel^−1^) for the second CBF3CCΔN dataset. The total exposure time of 10 s fractionated into 25 frames, a total dose of 46 e^-^ Å ^−2^ per movie for first dataset and the total exposure time of 15 s fractionated into 40 frames, a total dose of 60.9 e^-^ Å ^−2^ per movie for second dataset. Movies were aligned using MotionCor2 (Zheng et al., 2017).

### Image processing and model building

CTF parameters were estimated using CTFFIND4 (Rohou and Grigorieff, 2015) and CTF correction and following image processing were performed using RELION 2.0 (Kimanius et al., 2016), unless otherwise noted. Resolution is reported using the gold-standard Fourier shell correlation (FSC) (0.143 criterion) as described (Rosenthal and Henderson, 2003; Scheres and Chen, 2012) and temperature factors were determined and applied automatically in RELION 2.0. A subset of the initial dataset was picked using an automatically generated Gaussian reference by Gautomatch (Urnavicius et al., 2015), extracted using a 200^2^ pixel box and then subjected to reference-free 2D classification. Some of resulted 2D class averages from different views were selected to be low-pass filtered to 25 Å and used as references for further automatic particle picking of the initial dataset. The automatically picked particles were screened manually followed by reference-free 2D classification, which yields 69,392 particles for subsequent processing. An ovoid generated from SPIDER (Shaikh et al., 2008) was used as an initial model for 3D classification. The best 3D class was used to perform a 3D auto-refinement against all the good particles, resulting a 4.9 Å map. After substitution of the particles contributing to this map by re-extraction from dose-weighted images calculated by MotionCor2, a further 3D auto-refinement provided a reconstruction at 4.7 Å overall resolution. 212,724 particles from the second data set were picked from dose-weighted images and selected for further processing after reference-free 2D classification. After joining the two dataset, 282,116 particles were input to 3D classification using an initial 3D reference obtained by low pass-filtering (50 Å) the reconstruction of 4.7 Å map. Two different conformations of complex were obtained after 3D classification, with 187, 606 particles in an “openn” conformation and 94, 510 particles in a “closed” conformation. 3D auto-refinement of the 3D classes against the corresponding particles resulted in reconstructions of 4.1 Å map for the “open” conformation and 4.5 Å map for the “closed” confirmation, respectively. The maps from refinement were post-processed by RELION and sharpened by a negative B-factor using an automated procedure resulting in a 3.7 Å reconstruction for the “open” confirmation and 4.1 Å reconstruction for the “closed” conformation. Local resolution was estimated using RELION. Cep3 (Purvis and Singleton, 2007) and the BTB/POZ domain of Skp1 (Orlicky et al., 2003) were placed in the map using Chimera. Ctf13 was built *de novo* using Coot (Emsley et al., 2010) in an early 4 Å map, utilizing secondary structure predictions from both Psipred (Jones, 1999) and Phyre2 (Mezulis et al., 2015), sequence conservation between common ascomycetes and with reference to structural preferences for leucine rich repeat domains (Bella et al., 2008).

Sequence alignments were generated using ClustalW (Larkin et al., 2007) and annotated using ESPript (Gouet et al., 2003). A single round of Real Space Refinement in Phenix was used to refine geometry (Afonine et al., 2012). Tracing of the main chain was assisted using a 6 Å-filtered map. This model was then rigid body fit into the final map at 3.7 Å followed by a final polish in Coot and refinement with a single iteration of Phenix using the Real Space Refinement protocol. Secondary structure restraints were initially generated using the Cep3 and Skp1 crystal structures and, for Ctf13, from within Phenix, with manual editing where deviations from the crystal structures was evident. In order to validate the model an FSCfree and FSCwork were calculated. Using Phenix, the atomic coordinates of the final model were randomly shifted by 0.5 Å and subsequently real space refined against one unmasked half map (the 'working' map). The resulting model was converted to a map and an FSC calculated between it and both the working map (FSCwork) and the free half map (FSCfree).

### Gel electrophoretic mobility shift binding assays (EMSA)

Protein-DNA interactions were evaluated by EMSA. 24 pmol doubly labeled 33 bp CDEIII dsDNA (AATATTAGTGTATTTGATTTCCGAAAGTTAAA) were mixed with different amounts of protein with the indicated ratio of DNA: protein in the reaction buffer (25 mM Hepes, pH8.0, 200 mM KCl, 2 mM DTT, 10% glycerol, 0.02% NP-40, 10 mM MgCl_2_, 10 µM ZnCl_2_). For competition EMSA, the unlabeled competitor DNAs were 50 times more concentrated than the labeled DNA. The mixtures were incubated at room temperature for 45 minutes and resolved on a 3%-12% Bis-Tris Native polyacrylamide gel at a constant voltage of 150 V at 4°C in 1x Native PAGE running buffer for 110 minutes. After electrophoresis, the gel was scanned using an FLA-3000 fluorescent image analyzer (Fujifilm) excited with a 490nm laser.

## Author contributions

Conceptualisation: C.K.V.; Methodology: C.K.V., W.Z. and N.L.; Investigation: W.Z., N.L. & S.M.; Writing-original draft: C.K.V. & W.Z.; Writing –review & editing: C.K.V., W.Z. & N.L.; Visualisation: C.K.V.; Supervision: C.K.V. & N.L.; Funding acquisition: C.K.V.

## Acknowledgements

We are grateful to: Martin Singleton for the Ctf13 and Cep3 genes; Kiyoshi Nagai for sharing his yeast expression system and helping us test expression of our constructs; Dan Clare, Corey Hecksel and Alistair Siebert for data collection at eBIC. W.Z. and C.K.V. were funded by the BBSRC.

